# Climatic Factors in Relation to Diarrhea for Informed Public Health Decision-Making: A Novel Methodological Approach

**DOI:** 10.1101/545046

**Authors:** Takayoshi Ikeda, Thandi Kapwata, Swadhin K. Behera, Noboru Minakawa, Masahiro Hashizume, Neville Sweijd, Angela Mathee, Caradee Yael Wright

## Abstract

**Background:** Diarrheal disease is one of the leading causes of morbidity and mortality globally, particularly in children under 5 years of age. Factors related to diarrheal disease incidence include infection, malnutrition, and exposure to contaminated water and food. Climate factors also contribute to diarrheal disease.

**Objectives:** We aimed to explore the relationship between temperature, precipitation and diarrhea case counts of hospital admissions among vulnerable communities living in a rural setting in South Africa.

**Methods:** We applied a novel approach of ‘contour analysis’ to visually examine simultaneous observations in frequencies of anomalously high and low diarrhea case counts occurring in a season and assigning colors to differences that were statistically significant based on chi-squared test results.

**Results:** There was a significantly positive difference between high and low ‘groups’ when there was a lack of rain (0 mm of cumulative rain) for 1 to 2 weeks in winter for children under 5.

Diarrhea prevalence was greater among children under 5 years when conditions were hotter than usual during winter and spring.

**Discussion:** Dry conditions may lead to increased water storage raising the risks of water contamination. Reduced use of water for personal hygiene and cleaning of outdoor pit latrines affect sanitation quality. Rural communities require adequate and uninterrupted water provision and healthcare providers should raise awareness about potential diarrheal risks especially during the dry season.

## Introduction

An estimated 3.4 million people die from diarrheal and other water-related diseases each year [1]. Diarrheal disease is responsible globally for 21% of deaths per year in children younger than 5 years of age [2] and is ranked as the third leading cause of death in this age group in South Africa [3]. The transmission of diarrheal disease is determined by factors related to, among others, weather variables, the vector and agent, socio-economic and ecological conditions, and intrinsic human immunity [4].

Several infectious water-borne diseases, including diarrheal disease, are linked to fluctuations in weather and climate [5] and usually exhibit typical seasonal patterns in which the role of temperature and rainfall has been documented [6] but not always in agreement. Both floods and droughts can increase the risk of diarrheal diseases, although the evidence for the effects of drought on diarrhea is inconclusive [7]. Major causes of diarrhea, i.e. cholera, cryptosporidium, *E. coli* infection, giardia, shigella, typhoid, and viruses such as hepatitis A, are associated with heavy rainfall and contaminated water supplies [8]. In the tropics, diarrheal diseases typically peak during the rainy season. A significant association of non-cholera diarrhea related hospital visits was found with high and low rainfall and with high temperature in Dhaka, Bangladesh [9]. In Senegal, there were two annual peaks in diarrheal incidence: one during the cold dry season and one during the rainy season. Thiam et al. [10] observed a positive association of diarrheal incidence with high average temperature of 36 °C and above, and high cumulative monthly rainfall at 57 mm and above. In Vietnam, considerable spatial heterogeneity existed in the risk of all-cause for diarrhoea across districts investigated with low elevation and differential responses to flooding and air temperature, and humidity drove further spatial heterogeneity in diarrheal disease risk [11]. In Ecuador, heavy rainfall events were associated with increased diarrhoea incidence following dry periods and decreased diarrhoea incidence following wet periods [12].

Children and especially children under 5 years of age, are particularly susceptible to diarrheal disease. In a study using Demographic and Health Survey data from 14 Sub-Saharan countries, regional prevalence of diarrhoea in children under three years of age was considered in relation to variations in precipitation and temperature [13]. Results showed that shortage of rainfall in the dry season increased the prevalence of diarrhoea across Sub-Saharan Africa. Such shortages occur in many regions when rainfall is average to below-average relative to the long-term monthly-mean. The results also showed that an increase in monthly-average maximum temperature raises the prevalence of diarrhoea while an increase in monthly minimum temperature reduces the number of diarrheal cases [13]. Maximum temperature and extreme rainfall days were also reported as strongly related to diarrhoea-associated morbidity with the impact of maximum temperature on diarrhoea-associated morbidity appearing primarily among children (0–14 years) and older adults (40–64 years), but with relatively less effects on adults (15–39 years) [14].

Since diarrheal disease is a major cause of morbidity and mortality, particularly among children under 5 years of age in developing countries and given that climate change-related health consequences of diarrheal diseases are projected to pose significant risks to future populations [15] we set out to explore the relationship between climate factors (temperature and precipitation) and diarrhoea prevalence among vulnerable rural communities in South Africa located in a subtropical setting, using a novel approach. In this study, we used “contours” to visualize frequencies of diarrhoea anomalies occurring in a season and assigned colors to differences that were statistically significant based on chi-squared test results. Meteorologists and oceanographers apply this technique called ‘contour analysis’ to visually explain simultaneous observations [16]. Isopleths, or lines of equal value, are used in contour analysis to link places of equal parameter. The use of “contours” or “contour plots” is seen in climate-related research, for instance, to consider weather patterns [17]. To the best of our knowledge, this is the first-time contour analysis has been used in climate-based-health research for public health decision-making. The results may be useful for integration into early warning systems that use climate and other relevant information towards prevention of diarrhoea.

## Methods

### Data

Handwritten hospital admission records for 1 January 2002 to 31 December 2016 were collected from two large, public hospitals, namely Nkhensani Hospital and Maphutha L. Malatjie Hospital, located in Mopani District Municipality in Limpopo Province, South Africa (Figure 1). Hospital records were scanned using an SV 600 overhead snap scanner, pages were saved as soft copies as .pdf files and later printed for double data entry into an electronic database using EpiData (version 3.1). Each hospital admission record included patient’s name and surname, patient’s residential address, patient’s date of birth, patient’s age, date of admission and reason for admission.

**Figure 1.**
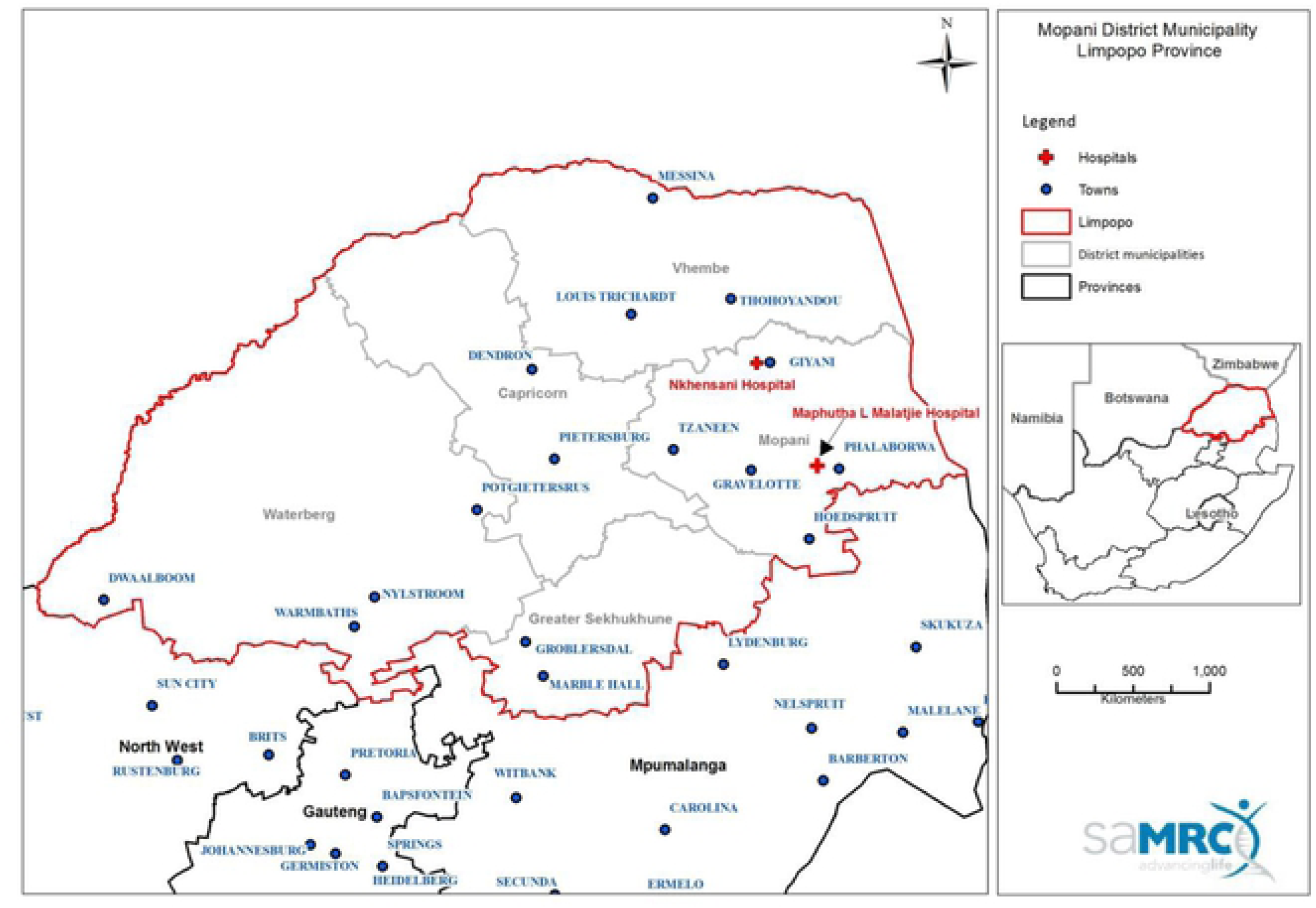
Location of the two hospitals in the study site in Limpopo Province, South Africa.

All diarrhea cases were extracted from the hospital admission records database for cases defined as diarrhoea using the criteria and terms provided by a South African medical doctor. Abdominal distention was not included since it could be associated with a variety of medical conditions other than diarrhoea. Data were unavailable in 2006 for both hospitals as well as at one hospital for weeks 1-23 in 2002 and weeks 1-40 in 2007. Despite the missing data mentioned above, our analyses could still be applied with the missing values (we did not replace missing values with zero) since we focused on anomalously high and low counts of daily admissions. It should be emphasized that the count for total admissions is not necessarily the total admissions at that hospital for that day / month / year but rather a total of the admissions that were captured by the hospital staff, collected by the researchers and entered by the data enterers. Cases of diarrhoea were summed as counts per week and diarrhoea weekly case counts were used in the contour analysis, described below.

Daily precipitation and temperature data were obtained from the South African Weather Service monitoring stations in the same District Municipality in which the two public hospitals were located. Table 1 presents the mean weekly precipitation and temperature by season for 2002 to 2016 for the study site. Precipitation data were available from one station in the District Municipality. Daily precipitation levels (mm) were summed to generate a weekly rainfall value. For temperature, data from 8 stations (namely Hoedspruit Air force Base, Tzaneen-Westfalia Estate, Levubu, Giyani, Tshivhasie Tea Venda, Tshanowa Primary School, Mukumbani Tea Estate and Punda Maria) at longitudes between 30.1 and 31.1° E and latitudes between 22.6 and 24.3° S in the District Municipality were extracted and applied in the study. Data from all these stations were used to calculate a spatially-averaged temperature for the study area. Daily minimum and maximum temperature (°C) values were then temporally averaged to generate weekly temperature values for minimum (Tmin) and maximum (Tmax) temperatures. For the contour analysis, weekly rainfall, Tmin and Tmax were used.

**Table 1.** Mean and standard error of weekly precipitation and temperature by season for 2002 to 2016 for the study site.

|  | **Season** |  |  |  |
| --- | --- | --- | --- | --- |
|  | **JJA** | **SON** | **DJF** | **MAM** |
| **Mean Tmax (° C)** | 24.0 | 28.7 | 29.7 | 26.9 |
| **Tmax SE** | 0.2 | 0.2 | 0.2 | 0.2 |
| **Mean Tmin (° C)** | 11.1 | 16.2 | 19.3 | 15.6 |
| **Tmin SE** | 0.1 | 0.1 | 0.1 | 0.2 |
| **Mean Prcp** | 0.6 | 7.4 | 19.5 | 6.9 |
| **Prcp SE** | 0.2 | 1.1 | 2.6 | 1.4 |

### Contour analysis

As an initial step for the analysis, we compared the climate variables, namely temperature and precipitation, between high and low diarrhea case count anomalies from hospital admissions, which were calculated by removing the climatological mean and linear trend of the case counts. The original time series was decomposed into three components: (i) Seasonal patterns that repeated with a fixed period of time (defined according to season as stated below) considered on a monthly scale and deemed as the climatological means; (ii) The underlying trend which could relate to the effort of collecting diarrhea data, the effect of some intervention or population growth, etc.; and (iii) The residuals of the original time series after the seasonal and trend series were removed referring to the “noise”, “irregular” or “remainder”, which is thus termed as the random component, i.e. the anomalies. Anomalies were therefore calculated according to Equation 1:

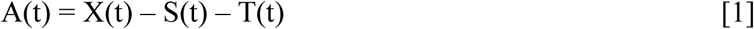

Where: A is the Anomaly, S is the seasonal component, T is the trend component, and t is time (or week). High weekly anomalies were inferred as ‘higher than normal’ diarrhea case counts and low anomalies were inferred as ‘lower than normal’, where ‘normal” refers to the long-term average for the corresponding period/week i.e. the first component discussed above. In addition, we discarded anomalies that were less than one standard deviation from the mean to retain anomalously extreme high or low incidence. Then we segregated the high and low incidence (case count anomalies) by season. Seasons were defined as spring: September-October-November (abbreviated as SON); summer: December-January-February (DJF); autumn: March-April-May (MAM); and winter: June-July-August (JJA).

For all weeks within each season, across the data set, we counted for precipitation how many times it rained on average ‘y’ mm over ‘x’ consecutive weeks (with x going from 1 to 10 weeks, and y for decile increments of the range of precipitation values) and tested these findings for statistically significant differences between the two groups of incidence category (i.e. high or low). The differences of how many times it rained in the high and low groups were plotted as contour lines. Differences with statistical significance, based on a chi-squared test for counts and using a Monte Carlo test [18] with 1 000 replicates to compute p-values at alpha level 0.01, were plotted. We repeated this approach for consecutive weeks with weekly Tmax and Tmin. For both precipitation and temperature, we also looked at lagged effects of each climate variable, for 0 to 8 weeks lag (chosen *a priori*) [9, 19], i.e. lag 3 means “temperature or rain 3 weeks prior to diarrhea counts of a certain week”. Contour plots were made for individuals of 5 years and older and for children under 5 years of age (i.e. 0 to 4 years), separately. All analyses were done in R version 3.2.2 [20]

### Ethics Statement

Permission to conduct the study was granted by the Limpopo Department of Health (REF 4/2/2), the management staff of Nkhensani Hospital and Maphutha L. Malatjie Hospital. Permission was granted by the South African Weather Service for use of the climate data. The South African Medical Research Council Research Ethics Committee approved the study protocol (EC005-3/2014).

## Results

### Hospital admission counts

Between 2002 and 2016 (inclusive) the total numbers (as captured for this study) of diarrhea hospital admission case counts at the two hospitals for individuals aged 5 years and older and for children under 5 years of age, separately, were 8 885 and 2 343, respectively. Figure 2 shows the high and low weekly diarrhea case count anomalies during the 14-year period for (a) individuals aged 5 years and older and (b) children under 5 years of age.

**Figure 2.**
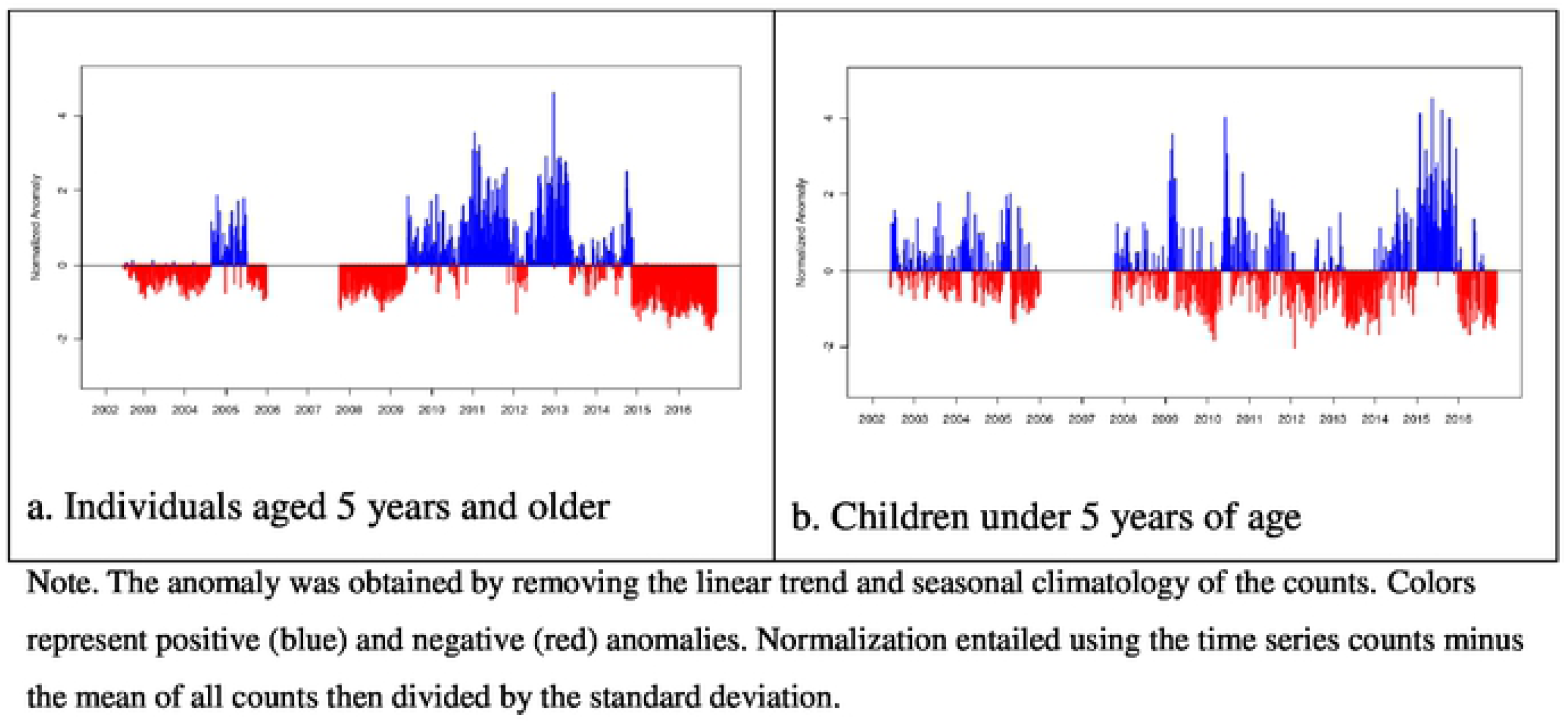
High and low weekly diarrhoea case count anomalies for (a) individuals aged 5 years and older and (b) children under 5 years of age between 2002 and 2016. Anomalies were normalized for comparison purposes.

### Precipitation and diarrhea case counts for individuals 5 years and older

We applied ‘contour analysis’ to the anomalously high and low weekly diarrhea count groups for individuals aged 5 years and older and compared these groups by counting consecutive weekly total rainfall. Figure 3 shows significant positive differences for different lags for JJA (dry season). At lags of 2, 3 and 6 weeks, cumulative rain of 8 – 14 mm for 6 to 10 consecutive weeks showed positive differences between high and low groups (orange colors). In the beginning of the rainy season SON, significant differences were seen in cumulative rain of up to 14 mm for 10 consecutive weeks for up to 2 weeks lag (Figure 4). In the rainy season (DJF), cumulative rain of 40 −52 mm for 8 to 9 consecutive weeks showed significant differences between high and low groups up to 1-week lag (Supplementary Figure S1). Similar levels of cumulative rain were seen in MAM, however, for lags of 5 to 8 weeks (Figure S2).

**Figure 3.**
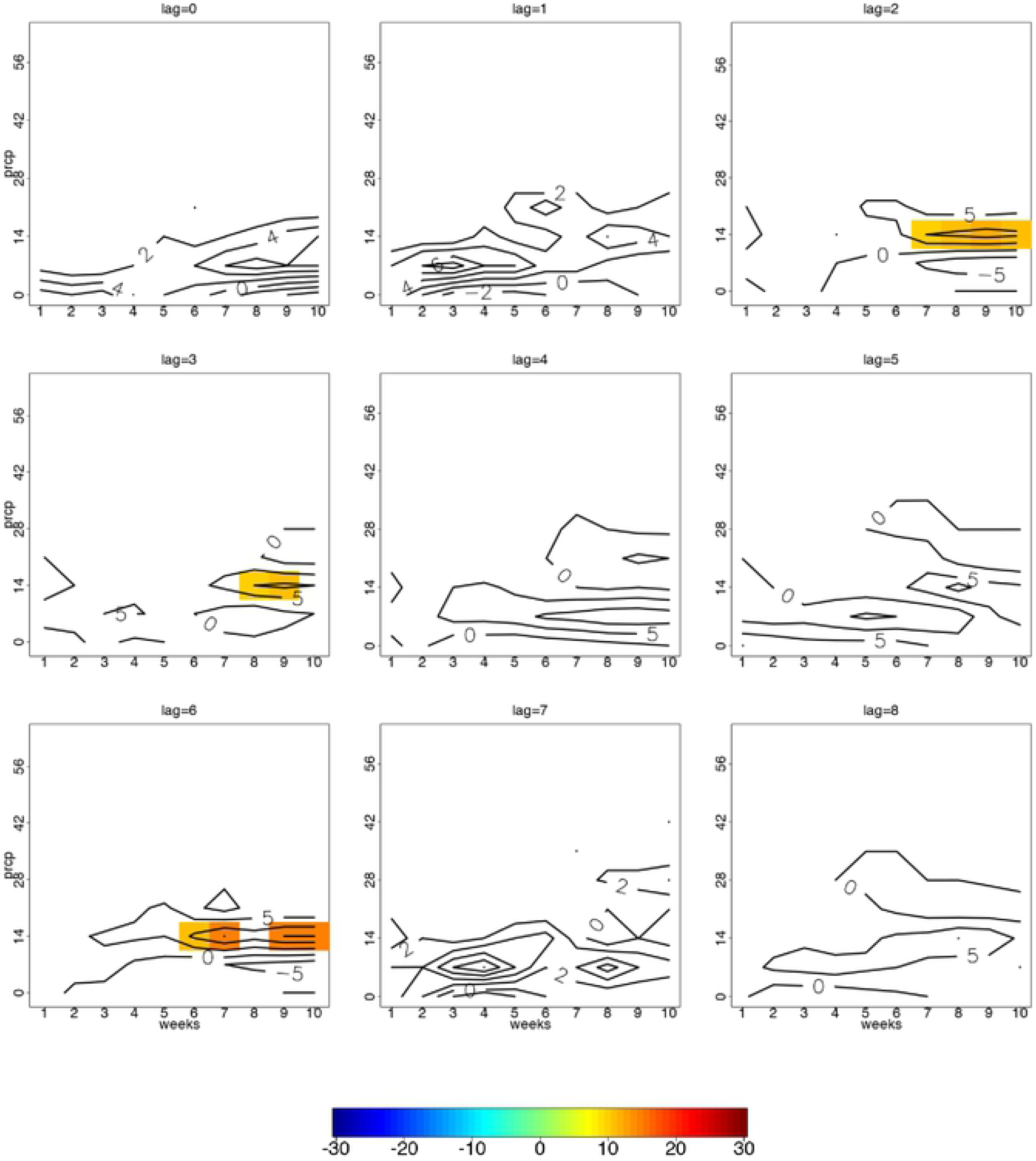
Statistically significant contour differences in anomalously high and low diarrhoea case counts for lag 0 to 8 weeks per consecutive weeks of precipitation among individuals aged 5 years and older for season JJA.

**Figure 4.**
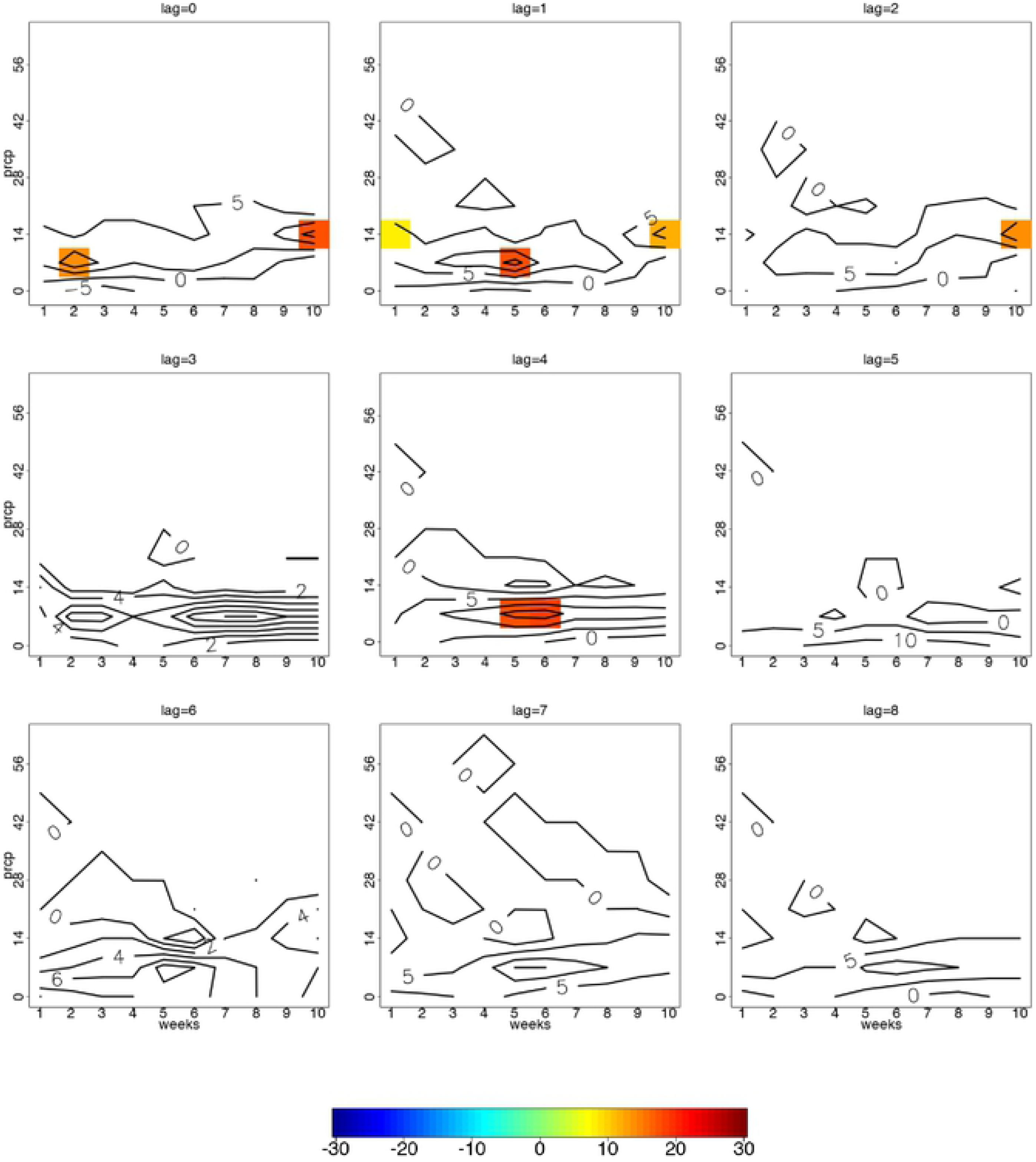
Statistically significant contour differences in anomalously high and low diarrhoea case counts for lag 0 to 8 weeks per consecutive weeks of precipitation among individuals aged 5 years and older for season SON.

### Precipitation and diarrhea case counts for children under 5 years of age

For children under 5 years of age, significant differences between the high and low groups showed different patterns in all seasons compared to the older age group. The most remarkable difference - evident by the ‘red cells’ in Figure 5 for most lags - indicated that there was a significantly positive difference when there was a lack of rain (0 mm of cumulative rain) for 1 to 2 weeks in JJA. Also, 5 or more consecutive weeks of 7 to 21 mm of cumulative rain showed significantly positive differences at most lags. For SON, significant differences were seen most noticeably at a lag of 5 weeks with 4 to 9 weeks of consecutively no rain (Figure 6). Significant differences were not seen in DJF (Figure S3) and MAM showed differences only at 4 and 8 for 8 to 10 weeks of consecutive cumulative rain of 14 – 26 mm (Figure S4).

**Figure 5.**
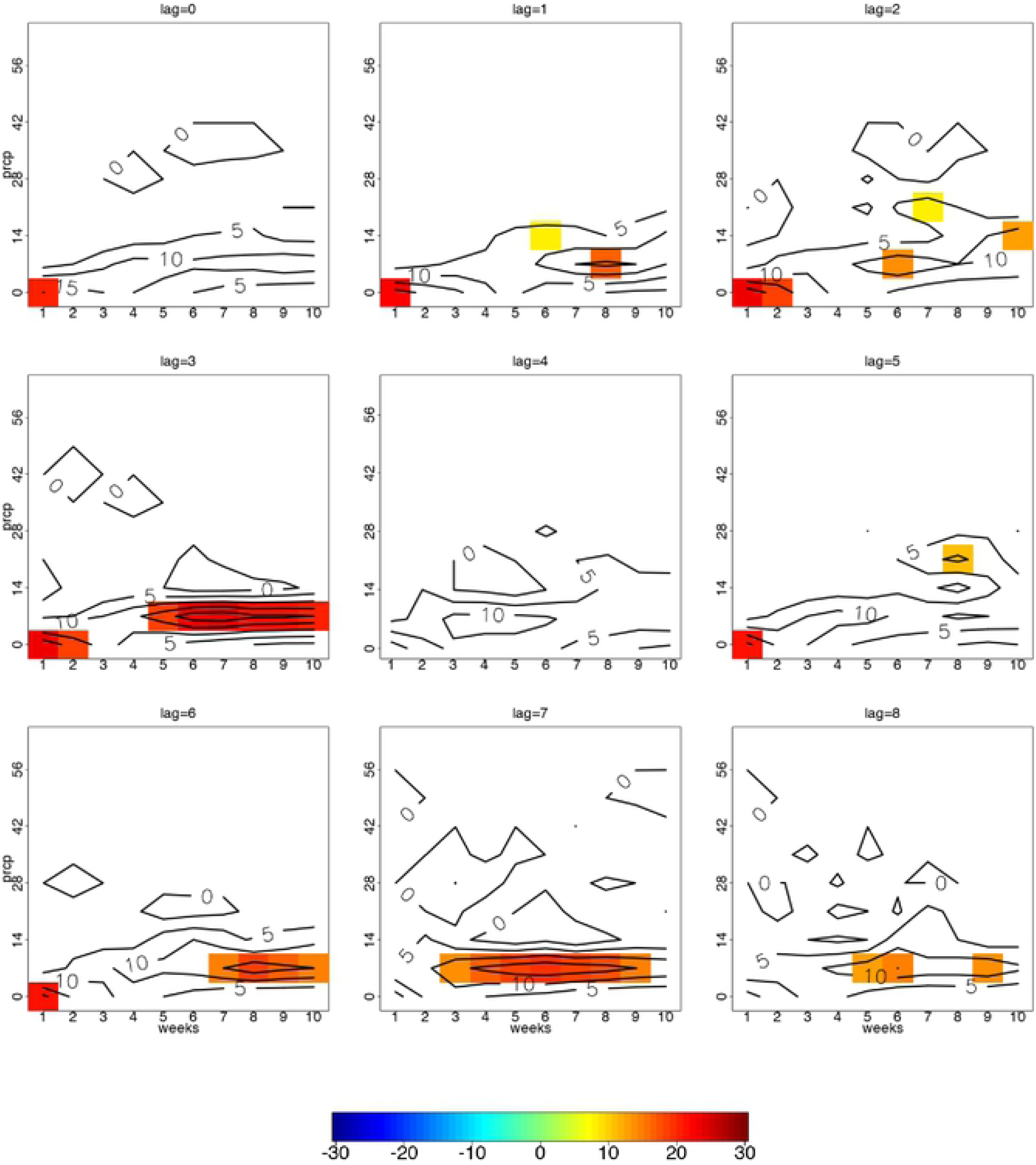
Statistically significant contour differences in anomalously high and low diarrhoea case counts for lag 0 to 8 weeks per consecutive weeks of precipitation among individuals under 5 years of age for season JJA.

**Figure 6.**
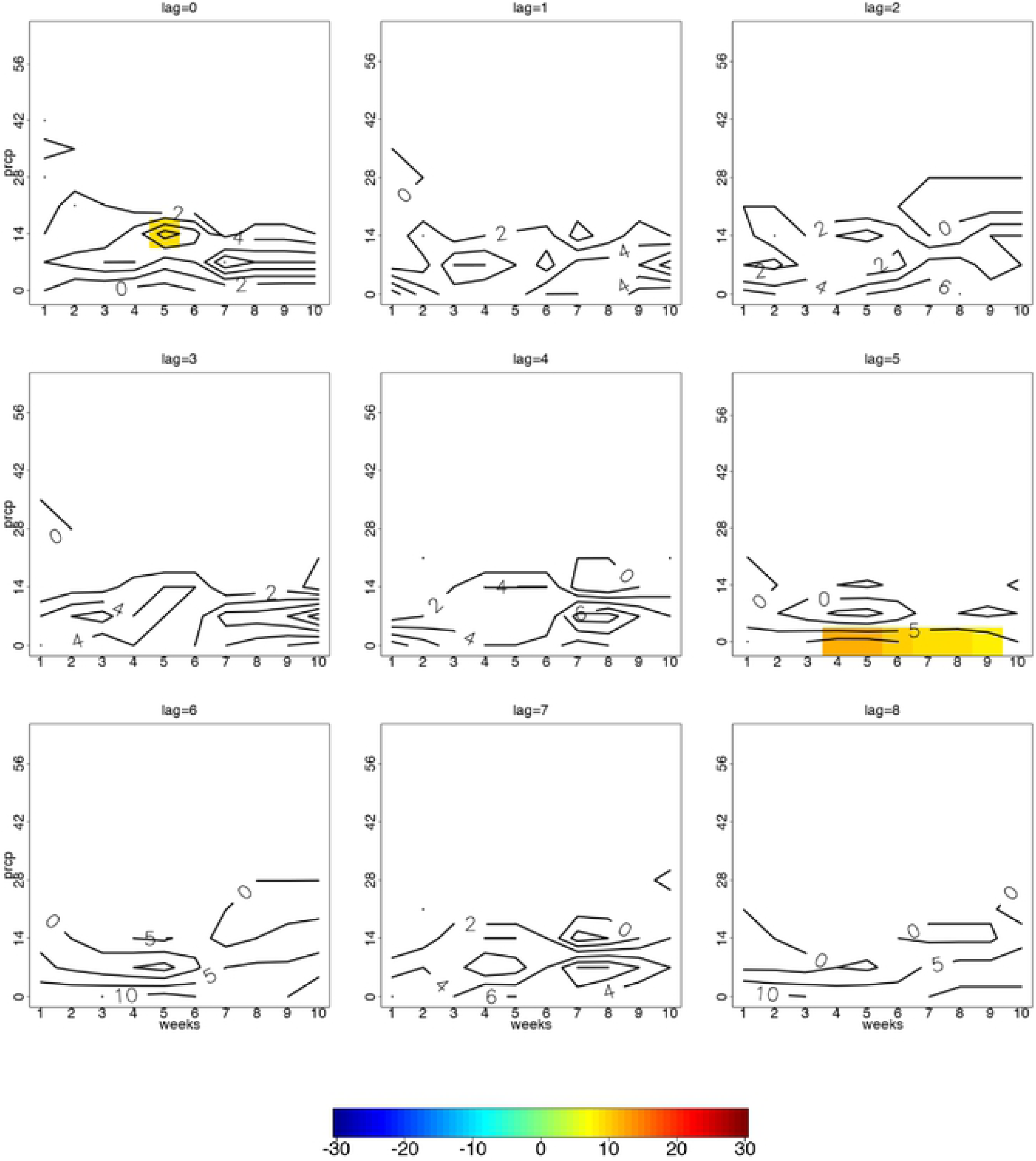
Statistically significant contour differences in anomalously high and low diarrhoea case counts for lag 0 to 8 weeks per consecutive weeks of precipitation among individuals under 5 years of age for season SON.

### Temperature and diarrhea case counts for individuals aged 5 years and older

There were no statistically significant associations between temperature (Tmin and Tmax) and high and low anomalies in case counts of diarrhea among individuals aged 5 years and older (Data not shown).

### Temperature and diarrhea case counts for children under 5 years of age

Among children under 5 years of age, there were some differences of Tmin and Tmax between the high and low groups for seasons JJA and SON. For JJA, 1 week of Tmin at 12 °C at 2 to 3 weeks lag and 16 °C at 8 weeks lag showed significant positive differences between high and low groups (Figure S5). For Tmax, 1 to 2 weeks of consecutive temperatures reaching 24 °C showed positive differences between the two groups at 3 to 4 weeks lag (Figure S6). As for SON, 9 to 10 weeks of consecutive Tmax of 26 °C showed positive significant differences at 5-, 7-and 8-weeks lag (Figure S7). Seasons DJF and MAM did not show any significant differences.

## Discussion

We set out to explore the relationship between precipitation / temperature and case counts of diarrhea hospital admissions using a novel approach of ‘contour analysis’ to visually explain simultaneous statistically significant observations in frequencies of anomalously high and low diarrhea case counts occurring in a season. Previous studies have used alternate statistical methods of analysis, such as time series regression [21, 22] to consider the relationship between precipitation and diarrhea, and temperature and diarrhea. In those studies, the datasets were significantly larger in size compared to the data available in our study and there were fewer missing data probably because of electronic record-keeping which is not common practice in rural, African hospitals and clinics. We successfully implemented contour analysis for the first time with meteorological and public health data. This method of analysis is therefore of great potential value for use with relatively small- and medium-sized datasets and where the hospital admissions data are constrained due to missing information - partially because of data being handwritten and not electronically captured. In time series analysis, for example, the approach would be to either remove all data prior to the complete year of missing data (in our study this was the year 2006) or use data imputation. In our analyses, we did not need to remove the data because our analysis was not affected by such missing data, instead, we treated missing values as missing (not zeroes) and focused on the anomalously high and low diarrhea counts.

Our most statistically significant findings were for children under 5 years of age in whom we saw a high prevalence of diarrhea when conditions were either wetter than average during the rainy season or drier than average during the dry season, as well as when temperatures were higher than normal. Children may be particularly vulnerable to diarrhea transmission when conditions are very dry and hot. A similar finding was seen in Nigeria where rotavirus was also most prevalent (95 % of all cases) during the dry season [23]. Similarly, in Botswana, diarrhea incidence is high in both the wet and dry seasons, but it was unexpectedly highest in the dry season with a 20 % increase over the yearly mean [24]. In the latter study, the authors hypothesized that the dry (and hot) conditions encouraged the activity and density of flies that transmitted diarrhea-causing microorganisms. We surmise that in Limpopo, South Africa, the warmer and drier conditions may lead to water shortages, lower availability of safe water sources [25]and increased water storage (perhaps not hygienically maintained), reducing personal use (and cleaning) of outdoor pit latrines, thereby reducing sanitation quality and personal hygiene. In addition, the wetter conditions may lead to increased risks of water contamination. Under-developed infrastructure and illegal connections to water supply pipes may also lead to contaminated water [26]. It is possible that children under the age of 5 are more vulnerable to such conditions. However, these assumptions remained to be verified among the communities served by the two hospitals from which data were drawn for use in the analyses presented here.

Based on the contour analysis results and associations found at certain lags, these results could assist healthcare practitioners to issue seasonal warnings of potentially high diarrhea and prevention measures in advance. For instance, separate warnings could be issued based on the contour analysis for individuals older than 5 years and for children under 5. For the younger age group, a warning could be issued for the winter season, up to 3 weeks in advance with up to 2 consecutive weeks of no rain. This warning can be updated on a weekly basis if there is still no rain occurring in the coming weeks. Most importantly, the warning would still be in effect if there is anomalous rain (7-21 mm) during the last 10 weeks prior to the winter season. Children under 5 were also susceptible to changes in temperature. For instance, when minimum temperature reaches 12 to 16 ° C, which is warmer than average for the winter season, it could also result in higher cases of diarrhea in the children under 5 years of age. It could be that wet and warmer conditions than average during winter co-vary and therefore anomalous winter rain may lead to warmer temperatures for that season and therefore increase diarrhea prevalence.

Our study was constrained by the quantity and quality of the hospital admissions data that were used to determine diarrhea prevalence. The medical records included those from the children, female and male wards only. All the records were handwritten and posed numerous challenges such as faded ink and handwriting being illegible, daily use of books leading to torn pages and errors in recording, for example, a male patient captured in a female ward. Misdiagnosis and misclassification were also evident in the hospital records, for example, where there was no clear indication of the final diagnosis of health outcome when a patient was discharged and where dates were entered incorrectly (the discharge date was sometimes captured as being before the admission date). While it would have been informative to know the cause of each diarrhea case, this information was not available from the hospitals, however, it is possible that the dry, cold season diarrhea was of viral cause [27]. In almost all cases the sex was missing. Therefore, it was not possible to use these data in our analyses. Despite those hurdles, the results presented here are statistically significant and missing and incorrect reporting are unlikely to drastically change the conclusions.

In summary, using a novel approach of analysis we detected trends in patterns of precipitation and temperature in relation to diarrhea prevalence for two separate age groups. Children under 5 years of age were especially vulnerable to diarrhea during very dry, hot conditions as well as when conditions were wetter than usual. We noted that local living, environmental and environmental health conditions may ‘overwhelm’ the typical climate-disease patterns known to influence diarrheal disease. Dry conditions lead to changes in water availability, use and storage and likely increase in the risk of diarrhea transmission in rural settings, whereas wet conditions lead to water contamination. Rural communities require adequate and uninterrupted water provision all-year round. Healthcare practitioners should help to raise awareness about potential diarrheal risks especially during the dry season. In addition to understanding climate-diarrhea patterns localized to specific settings and conveyed through early warning systems, a significant proportion of diarrheal disease can be prevented through safe drinking water and adequate sanitation and personal and food hygiene [1] which should remain a priority especially in rural settings. Finally, there is an opportunity to use contour analysis in public health and other sectors to improve planning and responses to, not only diarrhea, but other infectious diseases.

## Acknowledgements

This research was carried out for the iDEWS (infectious Diseases Early-Warning System) project supported by SATREPS (Science and Technology Research Partnership for Sustainable Development) Program of JICA (JAPAN International Cooperation Agency)/AMED (Japan Agency for Medical Research and Development) in Japan and the ACCESS (Applied Centre for Climate and Earth Systems Science) program of NRF (National Research Foundation) and DST (Department of Science and Technology in South Africa). Zamantimande Kunene, Mirriam Mogotsi and Wellington Siziba are acknowledged for collecting hospital data. We thank the hospital management and staff for permitting the study and for accommodating our team during the collection periods. We acknowledge the South African Weather Service for provision of the temperature and precipitation data.

